# Replication timing shapes the cancer epigenome and the nature of chromosomal rearrangements

**DOI:** 10.1101/251280

**Authors:** Qian Du, Saul A. Bert, Nicola J. Armstrong, C. Elizabeth Caldon, Jenny Z. Song, Shalima S. Nair, Cathryn M. Gould, Phuc Loi Luu, Amanda Khoury, Wenjia Qu, Elena Zotenko, Clare Stirzaker, Susan J. Clark

**Author notes:** Equal first authors (Qian Du, Saul A. Bert). **Corresponding Author**: Prof Susan J. Clark, Epigenetic Research Program, The Garvan Institute of Medical Research, 384 Victoria Street, Darlinghurst NSW 2010, Australia.

## Abstract

**Highlights:**

- Replication timing alterations are conserved in cancers of different cell origins
- Long-range epigenetic deregulation in cancer involves altered replication timing
- Cancer late-replicating loci are hypomethylated and acquire facultative heterochromatin
- Replication timing status potentiates *cis* and *trans* chromosomal rearrangements

**Summary:** Replication timing is known to facilitate the establishment of epigenome, however, the intimate connection between DNA replication timing and changes to the genome and epigenome in cancer remain uncharted. Here, we perform Repli-Seq and integrated epigenome analysis and show that early-replicating loci are predisposed to hypermethylation and late-replicating loci to hypomethylation, enrichment of H3K27me3 and concomitant loss of H3K9me3. We find that altered replication timing domains correspond to long-range epigenetically deregulated regions in prostate cancer, and a subset of these domains are remarkably conserved across cancers from different tissue origins. Analyses of 214 prostate and 35 breast cancer genomes reveal that late-replicating DNA is prone to *cis* and early-replicating DNA to *trans* chromosomal rearrangements. We propose that differences in epigenetic deregulation related to spatial and temporal positioning between early and late replication potentiate the landscape of chromosomal rearrangements in cancer.

## Introduction

Replication of the mammalian genome is an essential process that guarantees the accurate copying of genetic information before cell division. The regulation of this process during the synthesis (S-phase) of the cell cycle is fundamental for genome stability. The DNA replication program in a cell is defined as the temporal sequence of locus replication events that occur during S-phase, from early to late (Hansen et al., 2010; Rhind and Gilbert, 2013; Woodfine et al., 2004). This cellular program has been shown to stratify many features of the genome and epigenome; including gene density, gene transcription, histone modifications, DNA methylation and 3 dimensional (3D) chromatin organisation (Aran et al., 2011; Dileep et al., 2015; Julienne et al., 2013; Pope et al., 2014; Rivera-Mulia et al., 2015; Suzuki et al., 2011). Generally, “active” and open chromatin regions are replicated early in S-phase and “repressed” and closed chromatin regions replicated late in S-phase (Rhind and Gilbert, 2013). Alterations in the genome and epigenome are defining features of cancer cells (Baylin and Jones, 2016; Dawson, 2017) many of which contribute to unrestrained proliferation (Hanahan and Weinberg, 2011). Each round of replication represents an opportunity for error, leading to the acquisition of mutations (De and Michor, 2011) and copy number aberrations (Hodgkinson et al., 2012; Schuster-Bockler and Lehner, 2012; Woo and Li, 2012) in the genome. Epigenetic maintenance factors are associated with the DNA replication machinery (Alabert and Groth, 2012) and therefore DNA replication could also represent a similar opportunity for epigenetic deregulation commonly observed in cancer cells. However, the relationship between replication timing and epigenome alterations in cancer and the impact of these alterations in shaping the genomic landscape of tumour cells has remained largely uncharted.

Studies of mouse embryonic stem cell differentiation show that re-organisation of the replication timing program is accompanied by a concomitant re-organisation of the epigenome across large domains (Hiratani et al., 2010; Hiratani et al., 2008), demonstrating that the replication timing program is malleable and is an organisational process normally used to coordinate epigenetic identity. As the replication timing program contributes to both epigenetic maintenance and identity, a disruption of this process could be a key cellular event in carcinogenesis. We and others have previously shown that epigenetic deregulation in cancer encompasses large domains of long-range epigenetic silencing (LRES) and activation (LREA) with coordinated gene expression, histone modification, DNA methylation changes and disruption of Topologically Associated Domains (TADs) over several kilobases to megabases (Bert et al., 2013; Coolen et al., 2010; Taberlay et al., 2016). Ryba *et al.* (2012) have also shown that up to 18% of the genome can change in replication timing in pediatric leukemia. Given the long-range domain level of epigenetic change observed in cancer and the role of replication timing in epigenetic control, we hypothesise that the nature of replication timing landscape directs the mode of epigenetic remodelling and genomic alterations observed in cancer. Here, using high-resolution genome-wide characterisation of normal and cancer cells, we investigate how the replication timing landscape is associated with the cancer-specific epigenome changes and chromosomal rearrangements observed in prostate and breast cancers.

## Results

### The replication-timing program is largely conserved in normal and cancer cells

To ascertain if there are changes in replication timing in normal and cancer prostate cells, we performed Repli-Seq (Hansen et al., 2010) in duplicate in normal prostate cells (PrEC) and prostate cancer (LNCaP) cells and validated the technique at known loci of early (BMP1) and late replication (DPPA2) (Figures S1A and S1B) (Ryba et al., 2011). Immunoprecipitated DNA was sequenced and the density of reads were calculated for each cell cycle fraction in 50 kb sliding windows at 1 kb intervals (Table S1). To correct for local copy number variation and sequencing bias, the percentage of enrichment found in each fraction for a given locus was calculated as the Percent Normalised Density Value (PNDV) (Figure S1D). The PNDV values for each sorted fraction were further condensed into a single Weighted Average (WA) value, denoting the time of replication for that locus (Figure S1D). Replicate WA values for LNCaP and PrEC are highly correlated (r^2^ values >0.99) (Figure S1E). The distributions of WA values are comparable to WA distributions in other normal and cancer Repli-Seq datasets (Figure S1F).

To examine the replication timing landscape in the normal and cancer cells we first plotted the WA value for all ~2.8 million mappable 1 kb bins (loci). We found that 94.3% of loci are remarkably conserved in the normal and cancer cells (Figure 1A) using a stringent WA difference of 25 (|∆WA| < 25) (See Methods, Figure S1G). Comparatively, only 5.7% of the genome showed a difference in replication timing; 3.2% of the genome replicated *later* in LNCaP compared to PrEC (∆WA < −25), and 2.5% replicated *earlier* in LNCaP compared to PrEC (∆WA > 25) (Figure 1A). Next, to identify domains of consecutive loci in the cancer genome where the time of replication is altered, we merged all loci within 50 kb that had a |∆WA| > 25. This process yielded 314 domains that replicated *later* in LNCaP compared to PrEC with an average size of 288 kb, and 244 *earlier* domains that had an average size of 304 kb. The LNCaP replication altered domains are distributed across the genome spanning all chromosomes (Figure S2). An efficient replication origin cluster is shaped as an inverted V in the PNDV signal, indicative of a bidirectional replication fork progressing from early in S-phase (tip of inverted V) towards late (bottom of inverted V) (Hyrien, 2015; Petryk et al., 2016). Exemplary *later* and *earlier* domains are shown in Figure 1B and notably are also located in regions of inverted V shapes for both PrEC or LNCaP cells, suggesting that domains are associated with a change in replication origin firing.

**Figure 1:**
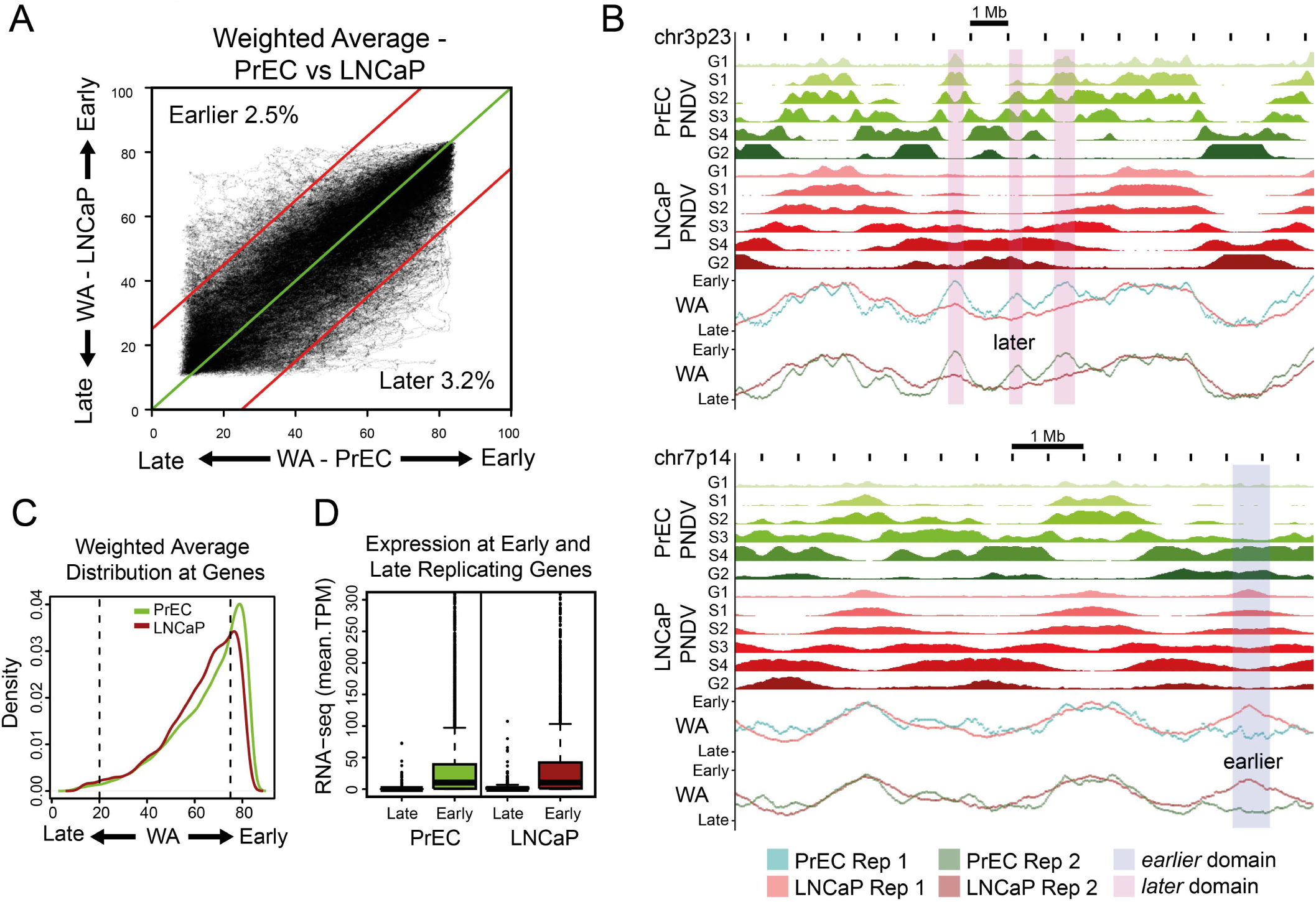
Replication timing is largely conserved between PrEC and LNCaP. (**A**) Replication timing weighted average (WA) values for all mappable 1kb loci are compared between PrEC and LNCaP. The green line indicates loci of no change in replication timing between PrEC and LNCaP. Loci outside the red lines (|∆WA| > 25) are substantially changed in replication timing. (**B**) Representative examples of regions of timing change between PrEC and LNCaP. PNDVs for PrEC (green) and LNCaP (red) are shown in the upper panels. The summarised WA values for each replicate are beneath. Regions that replicate *later* in LNCaP (∆WA < −25) are highlighted in pink. Regions that replicate *earlier* in LNCaP (∆WA > 25) are highlighted in blue. (**C**) The density of WA scores at gene promoters for PrEC and LNCaP. (**D**) Expression levels for genes that replicate early (WA > 75) and late (WA < 20) in PrEC and LNCaP.

### Differences in heterochromatin between normal and cancer cells occur predominately in late-replication

To investigate where replication timing stratifies the genome and epigenome landscapes, we first examined the genetic and epigenetic differences between PrEC and LNCaP at early- and late-replicating loci. We find that both PrEC and LNCaP cells display high gene density in early-replicating loci, but display low gene density in late-replicating loci (Figure 1C). Early-replicating genes tend towards higher expression and display higher plasticity. In contrast, late-replicating genes show constitutively low gene expression (Figure 1D). Next, we examined the genome-wide relationship between replication timing and chromatin features by performing a log odds ratio test. As expected from the expression data, we see positive associations between early replication and active marks and positive associations between late replication and repressive marks, in both cell lines (Figure 2A). In both PrEC and LNCaP cells, we find that active and permissive chromatin marks, including H3K4me3, H3K27ac, H3K4me1, H2AZac and H3K36me3 and DNAse1 hypersensitivity (HS), are progressively enriched towards early-replicating loci, whereas the repressive chromatin marks, H3K9me3, lamin A/C and lamin B1, are progressively enriched towards late-replicating loci in both cell types (Figures 2B, S3A and S3B). The relative difference in active and repressive chromatin enrichment with replication timing was also found in normal breast epithelial cell line (HMEC) and breast cancer cell line (MCF7) (Figures 2C, S3C and 2D).

**Figure 2:**
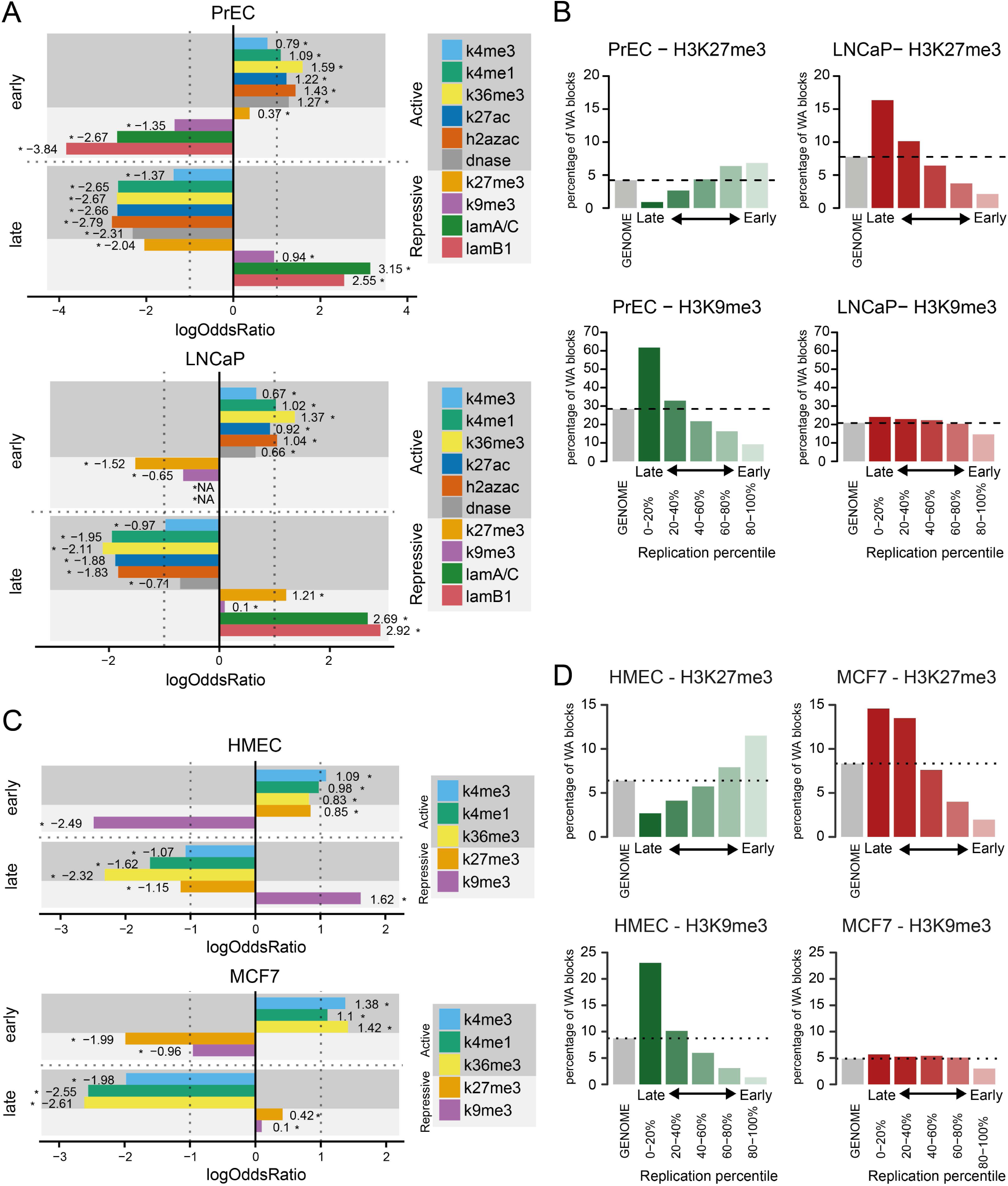
The relationship between chromatin and replication timing between normal and cancer cells. (**A**) logOdds ratio associations between histone marks, and early and late replication timing in PrEC and LNCaP. Association is above zero, and disassociation is below zero. Asterisks indicate significant associations (FDR < 0.05). (**B**) Percentage occupancy of chromatin marks for 1kb loci within replication timing percentiles for repressive marks H3K27me3 and H3K9me3 in PrEC and LNCaP. (**C**) logOdds ratio associations between histone marks in HMEC and MCF7, and early and late replication timing in MCF7. Association is above zero, and disassociation is below zero. Asterisks indicate significant associations (FDR < 0.05). (**D**) Percentage occupancy of chromatin marks for 1kb loci within replication timing percentiles for repressive marks H3K27me3 and H3K9me3 in HMEC and MCF7.

Interestingly, we identified two chromatin features, H3K27me3 and H3K9me3, which display distinct relationships with replication timing and differences in normal and cancer cells. First, H3K27me3 is more enriched in early-replicating loci in PrEC (6.82%), whereas in LNCaP, H3K27me3 is more enriched in late-replicating loci (16.34%) (Figure 2B). Second, H3K9me3 is enriched in PrEC late-replicating loci (61.77%) compared to early-replicating loci (9.21%), but the enrichment in late replication is reduced in LNCaP (Figure 2B). Opposing H3K27me3 and H3K9me3 enrichment at late-replicating loci are also seen in a comparison between HMEC and MCF7 (Figure 2D) suggesting that late-replicating loci is constitutive heterochromatin in normal cells and is more open facultative heterochromatin in cancer cells. Both polycomb and H3K9me3 remodelling is known to occur in cancer (Bender et al., 2013; McDonald et al., 2017; Pellakuru et al., 2012), but an inverse relationship had not previously been associated with replication timing.

Next we investigated whether the chromatin landscape is altered within the 5.7% of the genome that we found was different in replication timing from normal to cancer, based on the definition of |∆WA| > 25. We analyzed the differential presence or absence of chromatin feature peaks between PrEC and LNCaP in *earlier* or *later* replicating 1kb replication timing loci and tested for association using a log odds ratio test. Generally, we find that LNCaP show an enrichment of permissive marks at *earlier* replicating loci and depletion of permissive marks at *later* replicating loci (Figure S3D). In contrast, LNCaP show a depletion of repressive marks in *earlier* replicating loci and an enrichment of repressive marks in *later* replicating loci (Figure S3E). Interestingly, the repressive mark H3K27me3 does not show the same trend of reciprocal association as other repressive marks, but only shows enrichment towards *later* loci in LNCaP compared to PrEC (Figure S3E). Therefore, genome-wide remodelling of H3K27me3 in cancer appears to specifically occur at late or *later* replicating loci.

### Cancer-related DNA hypomethylation preferentially occurs in late replication

To next investigate the relationship between replication timing and DNA methylation we performed whole genome bisulfite sequencing (WGBS) in PrEC and LNCaP cells. Early-replicating loci in both PrEC and LNCaP cells display the highest levels of methylation (Figure 3A). Late-replicating loci in the normal cells have reduced DNA methylation levels compared to early-replicating loci. By contrast, there is dramatic reduction of methylation levels at late-replicating loci in LNCaP cells, indicating that cancer hypomethylation occurs preferentially at regions of late replication (Figure 3A). The dramatic decrease in methylation from early to late replication in LNCaP occurs at exons, introns, 3’UTR and intergenic regions, but not at CpG-island promoters as these are already predominantly unmethylated (Figure S4A). The extreme difference in methylation at late replication is also observed between breast normal HMEC and MCF7 cancer cells (Figure S4B). To further examine how replication timing relates to the overall methylation differences between the normal and cancer cells, we divided replication loci into hypomethylated (∆mCG < −0.2) and hypermethylated (∆mCG > 0.2) loci. First, we find that there are more hypomethylated than hypermethylated loci genome-wide (Figure 3B). Second, we find that hypomethylation is significantly more enriched in late-replicating loci compared to early replicating loci (74% vs 28%, p < 2.2e-16). In contrast, hypermethylation is significantly enriched in early-replicating loci compared to late-replicating loci (17% vs 2%, p < 2.2e-16) (Figure 3B). To further investigate the association between replication timing and DNA methylation, we visualized the data genome-wide and observe that on a broad scale, methylation is reduced in LNCaP compared to PrEC at regions of conserved late replication in both cell lines (Figures 3C and S4C). Strikingly, we find that unlike in PrEC where the DNA methylation and replication timing profiles are largely distinct, the genome-wide DNA methylation profile of LNCaP is more in synchrony with the profile of replication timing (Figures 3C and S4C). The correlation between DNA methylation and replication timing values increases from PrEC (Pearson’s 0.2213, Spearman’s 0.3073, p-value < 2.2e-16) to LNCaP (Pearson’s 0.5077, Spearman’s 0.4985, p-value < 2.2e-16). Together, the data demonstrate that late-replicating loci are predisposed to hypomethylation and early-replicating loci are predisposed to hypermethylation in cancer cells.

**Figure 3:**
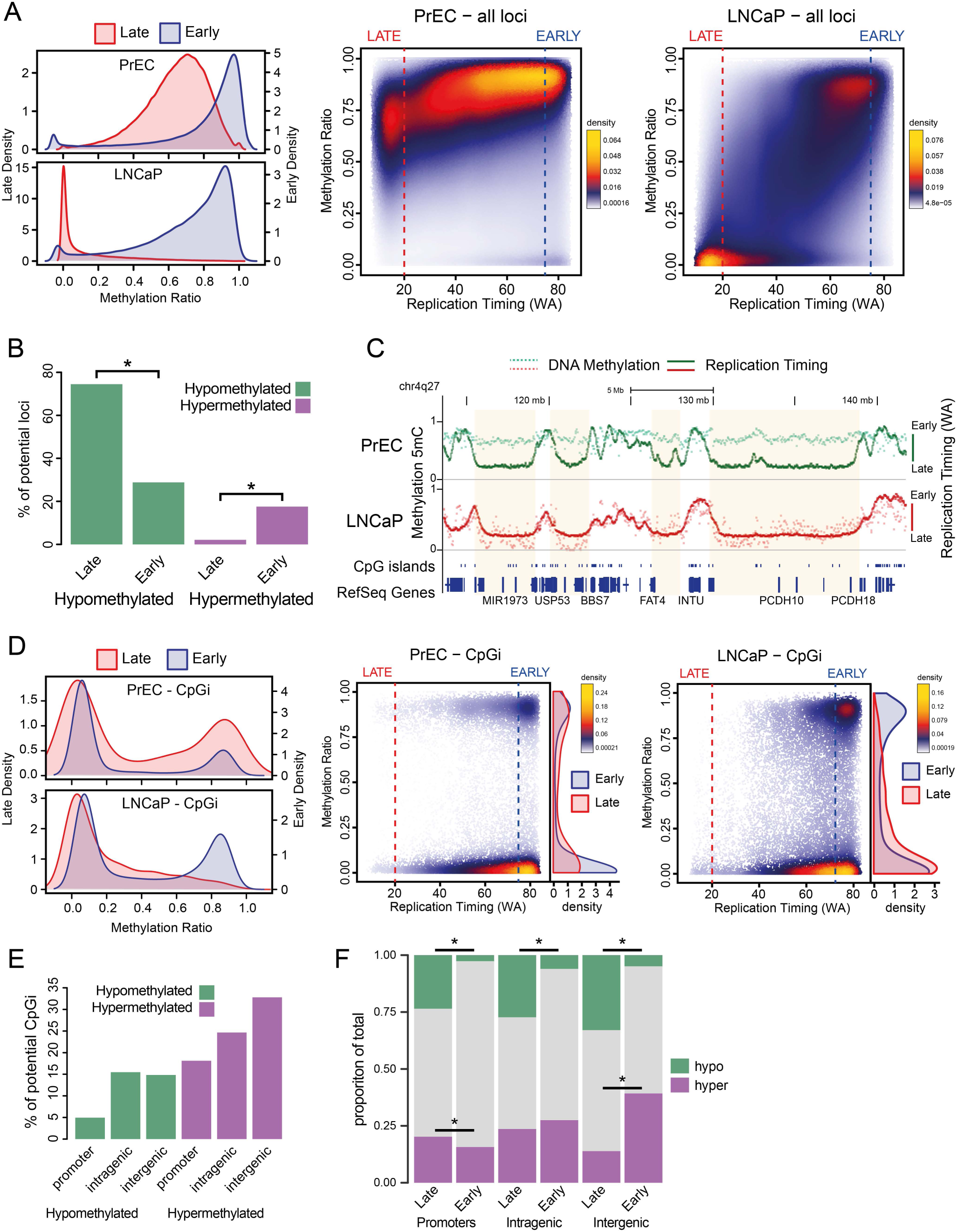
Replication timing correlates with DNA methylation in cancer. (**A**) DNA methylation (WGBS) density distributions for early (blue) and late (red) loci per cell line. Adjacent are scatterplots of DNA methylation in relation to replication timing (WA) for all measured 1kb loci in PrEC and LNCaP. Blue dashed line indicates early (WA > 75) and red dashed line indicates late (WA < 20). (**B**) Percentage of hypermethylated (∆WGBS > 0.2) and hypomethylated (∆WGBS < −0.2) 1kb loci in PrEC early and late regions. Asterisks indicate significantly different percentages between early and late (prop.test, p < 2.2e-16). (**C**) Representative examples of late-replicating regions common to both PrEC and LNCaP (shaded) that become hypomethylated in LNCaP. (**D**) DNA methylation (WGBS) density distributions for early (blue) and late (red) CpG-islands per cell line. Adjacent are scatterplots of DNA methylation in relation to replication timing for all CpG-islands. (**E**) Percentage of CpG-islands of each type that are hypermethylated or hypomethylated in LNCaP relative to PrEC. (**F**) Proportional plots of early and late CpG-islands that are hypo- or hypermethylated. Asterisks indicate significantly different percentages between early and late (prop.test, p < 0.05).

### Cancer-related promoter CpG-island hypermethylation preferentially occurs in late replication

We next investigated the relationship specifically between *CpG-island* methylation and replication timing. In PrEC, we found a bimodal distribution of methylation at both early- and late-replicating CpG-islands (Figure 3D). However, in LNCaP the bimodal distribution is only preserved in early-replicating CpG-islands, as late-replicating CpG-islands are predominantly unmethylated (Figure 3D). In addition, between the normal and cancer cells, we found that CpG-islands are predominantly hypermethylated (18.13%) in comparison to hypomethylated (4.95%) (Figure 3E). Interestingly, intergenic and intragenic CpG-islands show higher proportions of both hypomethylation and hypermethylation in LNCaP compared to promoters (Figure 3E), indicating that intergenic and intragenic CpG-islands are the most susceptible to epigenetic deregulation in cancer, rather than promoter CpG-islands. Importantly, we find that cancer-specific hypomethylated CpG-islands are enriched for late replication regardless of their location in promoters, intragenic or intergenic regions (Figure 3F). Similarly, hypermethylated promoter CpG-islands are enriched for late replication, whereas hypermethylated intergenic CpG-islands are enriched for early replication.

### Correlation of gene expression with replication timing and promoter methylation is dependent on CpG island status

DNA methylation and replication timing are both associated with gene expression, however it is unclear if there is an interdependency that defines gene activity. We therefore investigated the association of CpG methylation (averaged +/− 1000 bp around each TSS) with replication timing and gene activity. In both LNCaP and PrEC, over 70% of early-replicating genes contain CpG-island promoters, in comparison to only ~40% of late-replicating genes. We find that high levels of gene expression are associated both with an early time of replication and the presence of an unmethylated CpG-island promoter in both PrEC and LNCaP cells (Figure 4A). In contrast, high levels of gene expression are not observed for CpG-island associated genes that are unmethylated and late-replicating, or genes that replicate early and have methylated CpG-island promoters. The corollary of this observation is that both DNA methylation and replication timing a concerted remodeling a concerted remodeling in phase to promote high expression of CpG-island promoter associated genes. For non-CpG-island promoter genes, where transcription is generally lower, we observe transcriptional plasticity at early-replicating loci in both PrEC and LNCaP cells, regardless of the methylation status of the promoter (Figures 4B and S4D). At late replication, non-CpG-island genes are predominately inactive, independent of the promoter methylation status (Figures 4B and S4D). These late-replicating non-CpG-island genes are generally methylated in PrEC and unmethylated in LNCaP (Figure 4B). Therefore, in lieu of promoter CpG-islands, transcriptional output correlates more strongly with time of replication compared to promoter DNA methylation level.

**Figure 4:**
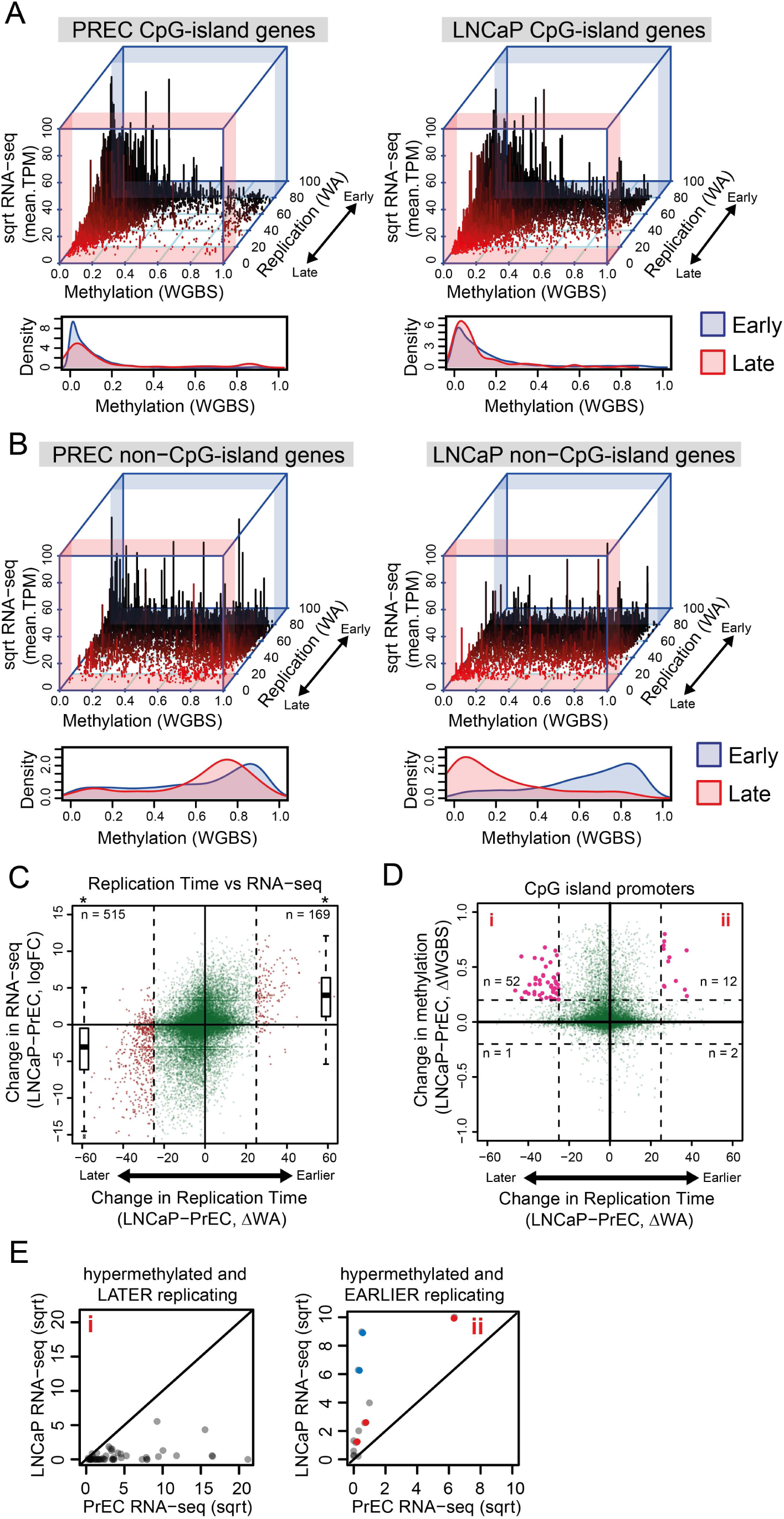
The relationship between replication timing, DNA methylation of gene promoters and expression. Replication timing is compared to DNA methylation and expression for CpG-island gene promoters (**A**) and non-island gene promoters (**B**) in PrEC and LNCaP. Underneath are methylation density distributions for early (blue) and late (red) promoters for each category. (**C**) Difference in replication timing (∆WA) compared to the change in gene expression (logFC). Dashed lines indicate a |∆WA| > 25, outside of which indicates a change in replication timing (red points). logFC boxplots for *later* or *earlier* genes are on the left and the right of the scatterplot, respectively. *Later* or *earlier* genes are significantly changed in expression compared to genes between |∆WA| > 25 (asterisk, Student’s T-test, p < 2.2e-16). - (**D**) Difference in replication timing compared to change in methylation at CpG-island promoters. Vertical dashed lines indicate a |∆WA| > 25 and horizontal dashed lines indicate a |∆WGBS| > 0.2. Promoters that are hypermethylated and either *earlier* or *later* are coloured in pink. Genes located in sections marked as **i** and **ii** are further explored in (**E**). (**E**) Comparison of expression levels (square root mean.TPM) between PrEC and LNCaP for hypermethylated and later gene promoters (**i**) and hypermethylated and earlier gene promoters (**ii**). Solid diagonal line indicates equal expression between PrEC and LNCaP; those above the line are more expressed in LNCaP and those below the line are less expressed in LNCaP. Highlighted are the two genes that are in Group I (blue), and the three genes that are in Group II (red) that had been previously defined (Bert et al., 2013).

### Differences in replication timing are associated with differences in the promoter epigenetic environment

We next investigated how a difference in replication timing in cancer is associated with gene expression within these regions, as well as the chromatin and methylation status. Using the definition of |∆WA| > 25, we find that more genes (515) are replicating *later* versus genes (169) that are replicating *earlier* in LNCaP compared to PrEC (Figure 4C). Genes that replicate *earlier* in LNCaP are significantly increased in expression, whereas the genes that replicate *later* are significantly repressed (Figure 4C). To investigate if replication timing changes of genes in cancer are also accompanied by chromatin remodelling, we compared the replication timing of genes to the enrichment of H3K27me3 and H3K4me3 at gene promoters (Figures S4E and S4F). We find that *earlier* replicating genes are associated with a more active promoter environment, characterized by concordant enrichment of H3K4me3 and depletion of H3K27me3. Similarly, we find that *later* replicating genes are associated with a more repressive promoter environment, characterized by concordant depletion of H3K4me3 and enrichment of H3K27me3.

We next asked if gene promoters that are located within regions of altered replication timing in cancer also display a change in DNA methylation. We find that there is distinct hypermethylation at promoter CpG-islands in cancer, regardless of the direction of replication timing change (Figure 4D). However, we do find that a higher proportion of CpG-island promoters become hypermethylated in *later* replication timing (n=52, 18%, Figure 4D Box i) relative to CpG island promoters that become hypermethylated in *earlier* replication timing (n=12, 11%, Figure 4D Box ii). We looked at the expression change of these hypermethylated genes and find that genes that replicate *earlier* in LNCaP tend towards increased expression (Figure 4Eii). By contrast, genes that are hypermethylated and replicate *later* in LNCaP tend towards decreased expression (Figure 4Ei). Interestingly, the hypermethylated and *earlier* replicating genes with increased expression in LNCaP compared to PrEC belong to two previously described groups of CpG-island promoter hypermethylation (Bert et al., 2013) where either, i) hypermethylation was associated with alternative promoter usage (*FRY, MCCC2*, and *CCDC67*, red circles, Figure 4Eii); or, ii) hypermethylation of the CpG-island borders resulted in augmentation of expression in prostate cancer (*NCAM and IQGAP2*, blue circles, Figure 4Eii). Together this data suggests that a change in replication timing is associated with a concerted change in both gene expression and the promoter epigenome.

### Late replicating regions associate with LADs and are regions of concordant DNA hypomethylation and heterochromatin change

Association with the nuclear periphery (nuclear lamina) is reported as a characteristic of late replication timing (Hansen et al., 2010; Peric-Hupkes et al., 2010; Pope et al., 2014), however, it is less clear how differences in nuclear lamina association relate to alterations in replication timing between cell types. To address whether differences in lamina association relate to differences in replication timing between normal and cancer, we performed ChIP-seq of both lamin A/C and lamin B1 in PrEC and LNCaP. Overall, we observe that LNCaP has 13% less lamina-bound DNA than PrEC, similar to others studies that show lamina association is reduced in cancer cells (Bell and Lammerding, 2016; Saarinen et al., 2015). We find that late replication timing is specifically characterized by the presence of both lamin A/C and lamin B1 rather than A/C or B1 alone (Figure 5A). Lamin A/C and lamin B1 lamina-associated domains (LADs) that are maintained from normal to cancer, show consistent late replication timing between normal and cancer, whereas LAD that are either ‘lost’ or ‘gained’ show a change in their replication timing program (Figure S5A). To examine if a change in LAD boundaries between normal and cancer also relates to replication timing change, we plotted a heatmap of WA values over the boundaries of PrEC LADs that are shifted in LNCaP (as shown by the white curve) (Figures 5B and S5B). These heatmaps demonstrate the remarkable correlation between borders of LADs and the transition from early timing outside the LAD to late timing within the LAD. A shift in the LAD boundary in LNCaP compared to PrEC correlates with a shift in where replication timing transitions from early to late, and ultimately where timing is different between PrEC and LNCaP (Figures 5B and S5B).

**Figure 5:**
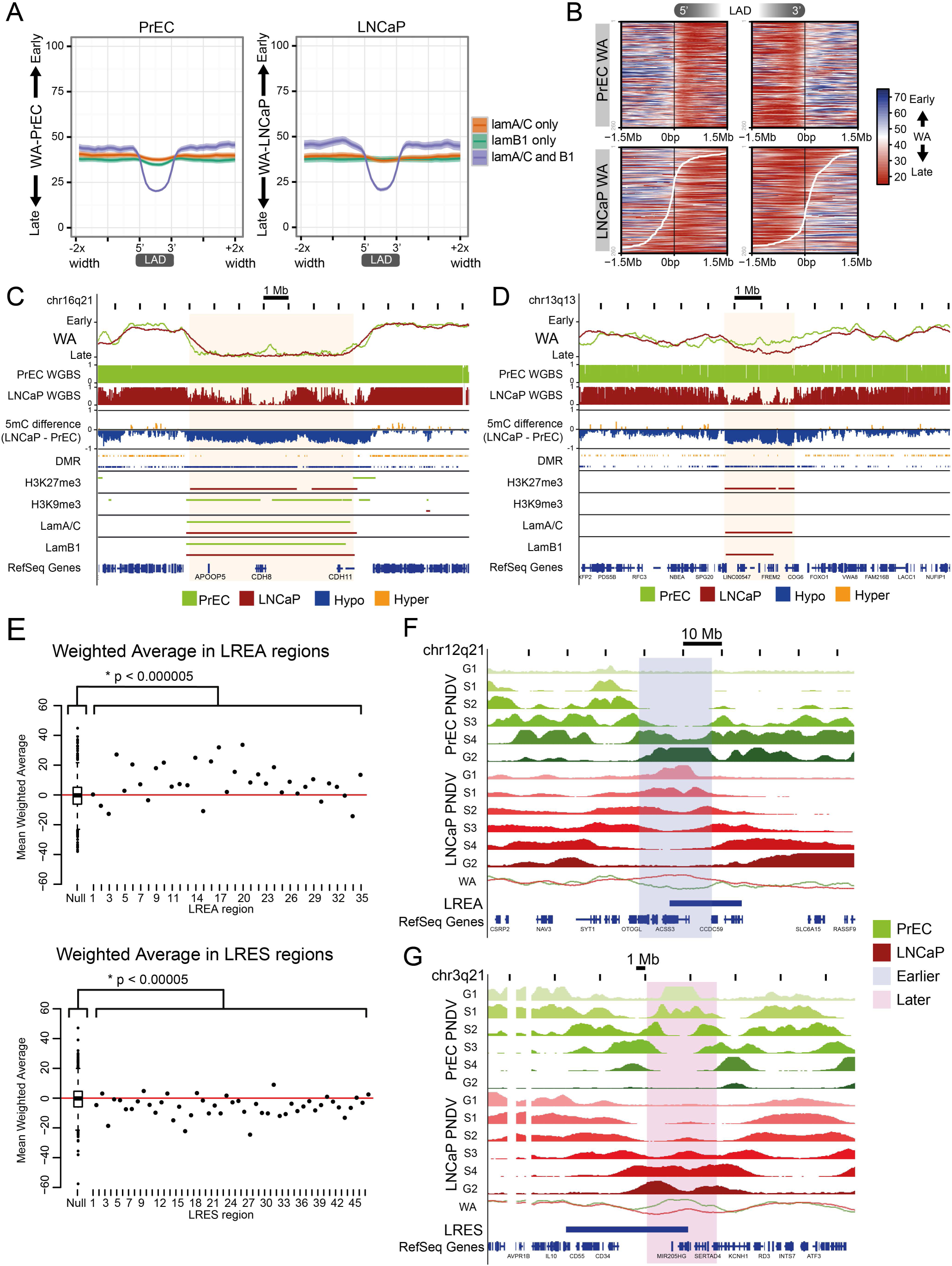
Long-range epigenetically regulated domains have altered replication timing in cancer. (**A**) Average plots of PrEC and LNCaP WA values over domains of lamin A/C-only, lamin B1-only or both. (**B**) Heatmap of PrEC and LNCaP WA values over the boundary of PrEC lamin B1 domains (LAD), ordered by degree of lamin B1 extension (upstream of 5’ and downstream of 3’) and loss (between 5’ and 3’) in LNCaP. Black lines down the centre of heatmaps represent PrEC LAD boundaries. White lines indicate the lamin B1 boundary in LNCaP. Scale for WA is from late (red) to early (blue). (**C**) Representative example of a late replicating region in LNCaP showing maintained LADs with coordinate DNA hypomethylation, H3K27me3 gain and H3K9me3 loss. (**D**) Representative example of a late-replicating region in LNCaP showing coordinate LAD gain, DNA hypomethylation and H3K27me3 gain. (**E**) The average replication time of each LREA or LRES region is compared to a distribution of 1000 randomized LREA or LRES regions (boxplot). The significance (Mann-Whitney test) of difference in WA distributions between regions and random is indicated. (**F,G**) Representative examples of overlaps between LRES and LREA and domains of replication timing (*earlier, later*).

Since we observed differences in both DNA methylation and repressive histone modifications between LNCaP and PrEC in late replication, we next investigated whether these epigenetic differences are located at nuclear lamina and co-occur at the same loci in LNCaP. Previous research has shown that DNA hypomethylation occurs in large domains that correspond to LADs or large domains of heterochromatin (Berman et al., 2012; Hon et al., 2012). Therefore, we called broad domains of H3K27me3 and H3K9me3 and annotated the 1kb replication timing loci for differences in the presence of histone domains, LADs and hypomethylated differentially methylated regions (DMRs), then tabulated the combinations (Figure S5C). Where the region is late-replicating in both PrEC and LNCaP, we find that the predominant chromatin pattern differences in LNCaP cells are for these regions to maintain lamina association from PrEC but to display concomitant DNA hypomethylation, H3K27me3 enrichment and either H3K9me3 depletion (17.40%, Figure 5C) or no presence of H3K9me3 (15.92%, Figure S5D). The next most common pattern shows acquisition of lamina association in LNCaP, a change to *later* replication and hypomethylation across the new LAD (6.9%, Figure 5D). Together our data shows that hypomethylation and heterochromatin alterations are occurring concordantly within the same late-replicating LAD in cancer.

### Replication timing changes comprise long-range epigenetically regulated domains

We have previously described large domains in the cancer genome which become coordinately epigenetically deregulated during tumourigenesis, termed either Long-Range Epigenetically Activated (LREA) or Silenced (LRES) domains (Bert et al., 2013; Coolen et al., 2010; Frigola et al., 2006). LREA (n=35) and LRES domains (n=47) were defined by coordinated changes in gene expression and histone marks. To further examine the relationship between replication timing and domains of concordant epigenetic change, we next asked whether LRES and LREA domains also change in replication timing. We find that LREA domains are significantly distributed towards *earlier* replication in LNCaP compared to PrEC and conversely, LRES domains are significantly distributed towards *later* replication (Figure 5E). By comparing to a random distribution of similarly sized regions, we find that LRES regions have a significant overlap with *later* domains (p=0.038) and LREA regions have a significant overlap with *earlier* domains (p=0.00097). Figure 5F highlights an exemplary LREA region (12q21) that is *earlier* in LNCaP compared to PrEC, forming an ectopic replication initiation zone. Conversely, Figure 5G displays a LRES region (3q21) that is *later* in LNCaP compared to PrEC, potentially due to loss of a replication initiation zone. More examples of LREA and LRES regions with differences in timing can be found in Figure S5E. In summary, long-range epigenomic transformation during tumourigenesis appears to be associated with a difference in replication timing together with a concerted remodelling of the epigenome to establish a cancer transcription program.

### Conservation of replication timing alterations in cancer

To further explore if there are similar patterns of replication timing differences across other cancers, we performed Principle Component Analysis (PCA) and hierarchical clustering using PrEC, LNCaP and publicly available ENCODE Repli-Seq data (Hansen et al., 2010; Thurman et al., 2007). These cell lineages include normal cultured primary cells, established fibroblast cell lines, Epstein-Barr Virus (EBV) transformed normal lymphoblastoid cells, embryonic stem cell (ESC) and five cancer cell lines HelaS3, MCF7, SK-N-SH, HepG2 and K562 (Figure 6A). In both PCA and hierarchical clustering, cell types were divided into 4 major clusters (Figures 6A and 6B). The normal cells are comprised of two clusters, one containing normal epithelial (PrEC), epidermal and endothelial cells, and the other fibroblasts. The normal cells are separate from the clusters containing the EBV transformed normal lymphoblasts and cancer cell lines. The closer clustering of EBV transformed normal lymphoblasts to cancer cell lines suggest the transformation process has indeed made these cells more cancer-like. Notably, all the cancer cell lines cluster together with the exception of HelaS3. The ESC line was also found to associate with the cancer group, potentially indicating a progenitor-like state of cancer. Since most cancer cells assayed cluster separately to normal cells regardless of cell of origin, this suggests that there may be a set of specific alterations in replication timing that are shared in cancer cells.

**Figure 6:**
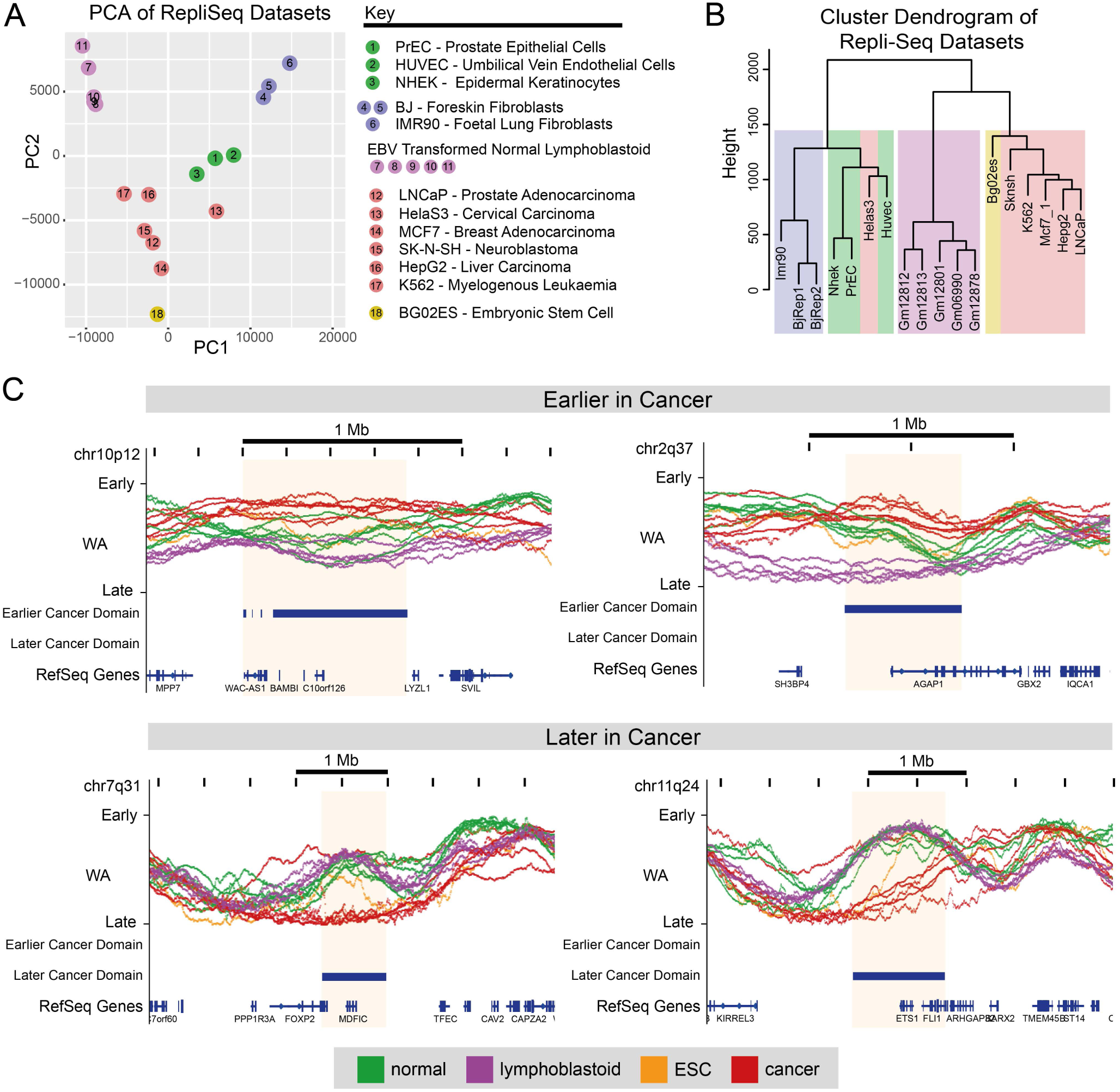
Conservation of replication timing alterations in cancer. WA values of all publically available Repli-Seq datasets, including PrEC and LNCaP, are assessed using PCA (**A**) and hierarchical clustering (**B**). Samples are identifiable by colour and number key. (**C**) Shown are representative examples of loci that are Earlier Cancer Domains (ECDs) or Later Cancer Domains (LCD) in multiple cancers compared to other cell types. Shaded boxes indicate ECDs or LCDs with logFC >= 1.

Next, we identified regions that displayed consistent differences in replication timing domains in the cancer datasets compared to all other non-cancer datasets. We identified 16 Earlier Cancer Domains (ECDs) and 56 Later Cancer Domains (LCDs). Representative examples of ECDs and LCDs across multiple cancer types are shown in Figure 6C. Interestingly, the majority of ECDs (15/16) and LCDs (55/56) overlap regions with apparent high timing variation, indicating these loci are innately malleable, yet notably distinct in timing in cancer compared to other cell types (Figure S6). Moreover, a subset of ECDs (10/16) and LCDs (30/56) are differentially timed between cancer and ESCs (Figures 6C and S6). Genes within ECDs and LCDs are given in Table S2 and include cancer-related genes such as ETS1, FOXP2 and BAMBI. Notably, 46.88% of ECD and 47.37% of LCD genes have ‘lincRNA’ status based on GENCODE 19 with LCD genes being significantly enriched in lincRNAs (prop.test, p = 0.00001128). To see if there is a potential indication of coordinate gene function, we relaxed our domain calling cutoff (See Methods) and looked for enrichment of GSEA terms. The analyses suggests that ECD genes may play a role in cell-to-cell adhesion and LCD genes may play a role in the cell’s immune response (Figure S7). The consistent differences in replication timing domains shared between cancer cells may therefore reflect the consolidation of a cancer-specific transcriptional program that occurs in all cancer transformations irrespective of tissue-of-origin.

### Relationship of replication timing with hypomethylation and genomic instability

It has been previously reported that DNA hypomethylation, heterochromatin remodeling and late replication timing predisposes the genome to chromosomal instability in tumourigenesis (Chen et al., 2007; De and Michor, 2011; Peters et al., 2001). To investigate if replication timing potentially influences the nature of chromosomal instability in prostate and breast tumourigenesis, we analysed publicly available clinical datasets (Baca et al., 2013; Berger et al., 2011; Robinson et al., 2015; Yang et al., 2013). We find in all prostate datasets that genomic rearrangement breakpoints are enriched in late-replicating DNA (WA < 20) and deplete in early-replicating DNA (WA > 75) in PrEC (Figure 7A). Chromosomal rearrangements were further classified as *cis* or *trans* depending on whether they were inter- or intra-chromosomal. We found that *trans* translocations are enriched at early-replicating loci in PrEC and *cis* rearrangements at late-replicating loci in PrEC in the Baca (2013) and Berger (2011) datasets (Figures 7B and 7C). We further divided the *cis* rearrangements into discrete subtypes and found inversions, deletions and long-range insertions were all enriched in late replication (Figure S8A). To further support this finding, we used published breast cancer WGS structural variation data from Yang *et al.* (2013) with MCF7 replication timing data. We also observe the same trend where *trans* variants are enriched in early-replicating loci and *cis* variants are enriched in late-replicating loci (Figures 7D and 7E). Taken together our data shows that the replication timing status of a locus can increase susceptibility to genomic rearrangement and notably, bias loci towards either *cis* or *trans* translocations.

**Figure 7:**
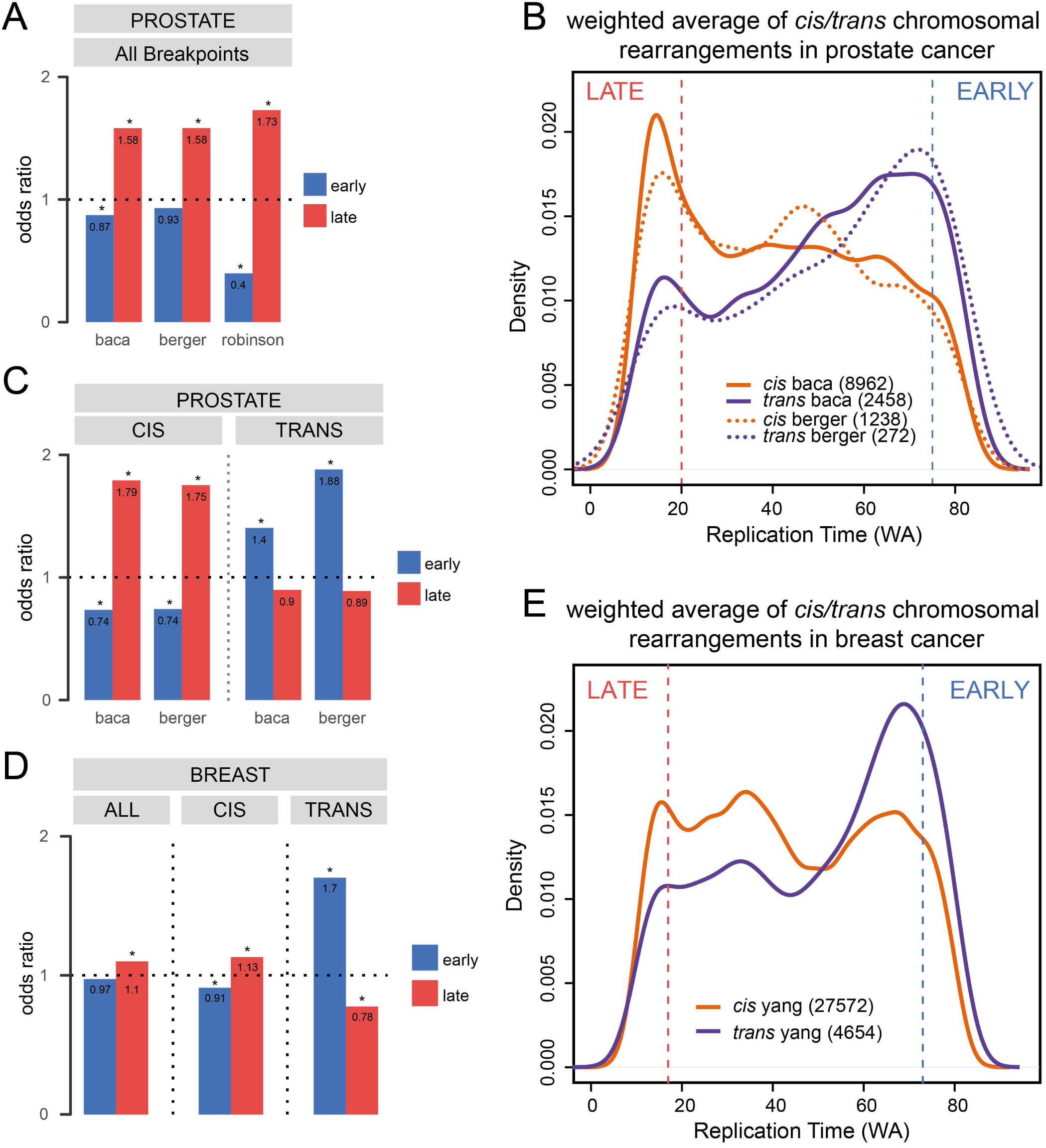
Replication timing stratifies the nature of chromosomal rearrangements in cancer. (**A**) Enrichment of rearrangement breakpoints for early or late replication in PrEC using public datasets Baca *et. al.* (2013), Berger *et. al.* (2011) and Robinson *et. al.* (2015). (**B**) Replication timing (PrEC WA) distributions for *trans* (purple) and *cis* (orange) rearrangement breakpoints. Solid lines are distributions for Baca *et. al.* (2013) data and dashed lines are distributions for Berger *et. al.* (2011). Vertical red and blue dotted lines indicate late (WA < 20) or early (WA > 75) cutoffs. (**C**) Enrichment of rearrangement breakpoints for early or late replication separated by *cis* or *trans* status. (**D**) Enrichment of rearrangement breakpoints all together or separated by *cis* or *trans* status for early or late replication in MCF7 using public dataset Yang *et al.* (2013). (**E**) Replication timing (MCF7 WA) distributions for *trans* (purple) and *cis* (orange) rearrangement breakpoints. Vertical red and blue dotted lines indicate late (WA < 17) or early (WA > 73) cutoffs.

We further investigated the nature of gene fusions that are commonly found in prostate cancer and asked if the genes were located in regions that displayed differences in replication timing. We examined replication timing states in PrEC and LNCaP of all the gene fusions documented in the Robinson (2015) dataset. Interestingly, we found that the majority of the breakpoints occurred in regions that shared the same state of replication timing (Figure S8B). Table S3 summarises all the breakpoints in prostate cancer including those that show significant replication timing differences. Of all gene fusions assayed, TMPRSS2-ERG located on chromosome 21q22.2 was the most common (209/4122) fusion found in patients. Strikingly, in 195/209 fusion events, the 3’ ERG translocation points were located in a domain that changes replication timing from early to late S-phase in LNCaP. Figure S8C shows the domain of replication timing change that harbors the ERG locus.

## Discussion

The major question we set out to address in this work was whether there are replication timing differences in cancer and if so what was the association between replication timing and alterations to the cancer genome and epigenome. Here, we made the remarkable discovery that replication timing is associated with the mode of chromosomal rearrangements in cancer due to the opposing remodelling of the epigenome in early versus late replication. We demonstrate that genomic regions that undergo long-range epigenetic deregulation in cancer also show concordant differences in replication timing. Notably late-replicating regions in prostate and breast cancer cells display a remarkable reduction of DNA methylation, and change in heterochromatin features from constitutive to facultative, which we propose results in a more open chromatin structure leading to an increase probability for chromosomal rearrangements. Moreover, our results illustrate that replication timing may play a critical role in determination of the location of both genetic and epigenetic alterations in malignancy.

The replication timing program is known to be re-organised during cellular differentiation and reflects cellular identity (Hiratani et al., 2010; Hiratani et al., 2008; Rivera-Mulia et al., 2015; Ryba et al., 2010). We find substantially fewer differences in prostate cancer replication timing compared to studies on ESC differentiation (~20-50%), but a similar percentage to the differences found between closely related somatic cells (4.5-6%) (Hansen et al., 2010; Hiratani et al., 2010; Hiratani et al., 2008; Ryba et al., 2010). This observation suggests that the basic mechanisms that control the replication program are not intrinsically deregulated in prostate tumourigenesis. However, despite the overall maintenance of replication timing between cell-types, it was intriguing to find that the cancer replication timing profiles are more similar to each other than to non-cancer cells. This suggests that a conserved alteration in replication timing exists in cancers of different origins, which may have potential functional relevance to tumourigenesis. We further identify domains that show similar alterations in replication timing between cancer and normal cells (ECDs and LCDs) and these domains consist of genes involved in cancer-related pathways, such as cell-to-cell adhesion and immunological signatures. It is therefore plausible that a change to *later* replicating timing may be involved in shutting down the immune response pathways in cancer cells. These ECDs and LCDs are also enriched in genes with lincRNA status suggesting that these regions may have a more regulatory role than protein coding role.

Domains of replication timing change we found in the prostate cancer cells correlate with previously observed domains of long-range epigenetic remodelling for both LREA and LRES regions (Bert et al., 2013; Coolen et al., 2010), whereby *earlier* domains become more ‘active’ and *later* domains become more ‘repressive’. This directional relationship has also been observed in development and differentiation (Hiratani et al., 2010; Hiratani et al., 2008; Rivera-Mulia et al., 2015), and further suggests that replication timing is a higher-order functional domain and a unit of epigenomic deregulation during tumourigenesis (Thurman et al., 2007). This higher-order organisation aligns with recent research showing the similarities between replication timing domains, units of higher-order chromatin organisation (TADs) and A/B compartments (Pope et al., 2014). Furthermore, the most widespread epigenetic alteration we observe between normal and cancer, in relation to replication timing, is DNA hypomethylation accompanied with a switch from constitutive to facultative heterochromatin within lamina-bound late-replicating regions. Previous research has reported several pairwise associations between late replication, lamina-associated domains (LADs), partially methylated and hypomethylated domains, and domains of H3K27me3 and H3K9me3 (Berman et al., 2012; Hon et al., 2012; Peric-Hupkes et al., 2010; Sadaie et al., 2013; Statham et al., 2012; Wen et al., 2009). However, our new findings indicate that there may be a highly co-ordinated alteration of the cancer epigenome that occurs exquisitely in genomic regions that replicate late in S phase. Of note, we observe a clear link between loss of H3K9me3, gain of H3K27me3 and DNA hypomethylation in late replication. The relationship between H3K27me3 and DNA methylation is generally considered to be antagonistic (Brinkman et al., 2012; Hon et al., 2012; Statham et al., 2012). In contrast to H3K27me3, H3K9me3 loss can contribute to DNA hypomethylation, as H3K9me3 is required to enhance the activity of UHRF1 and consequently DNMT1 (Liu et al., 2013; Rothbart et al., 2012; von Meyenn et al., 2016).

Finally, we observe an association between chromosomal rearrangements and replication timing, with later replicating regions prone to *cis* rearrangements and early-regions to *trans* rearrangements. Although associations between replication timing and chromosomal rearrangements have been previously been reported, the nature of the translocations was not deduced (De and Michor, 2011; Shugay et al., 2012). Our new combinatorial epigenome and replication timing data therefore leads us to propose a new model for how replication timing status and associated epigenetic alterations may influence the nature of chromosomal rearrangements (Figure 8). Interphase chromosomes are organized such that early-replicating loci are gene-dense, transcriptionally-permissive, ‘open’ chromatin, located towards the nuclear center and often close in proximity with other chromosomes (Bickmore, 2013; Gilbert, 2002; Goldman et al., 1984; Yaffe and Tanay, 2011). Late-replicating loci are gene-poor, transcriptionally inactive, condensed chromatin, located towards the nuclear periphery (lamina-bound) and typically contained within themselves (Bickmore, 2013; Gilbert, 2002; Goldman et al., 1984; Yaffe and Tanay, 2011). These typical patterns are reflected in the normal prostate cells (Figure 8). However we find that late-replicating loci in cancer are hypomethylated and become de-condensed to a ‘looser’ heterochromatin state. Even though many of these changes have been separately linked to increased genomic instability in cancer (Branco and Pombo, 2006; Chen et al., 2007; Kondo et al., 2008; Peters et al., 2001; Putiri and Robertson, 2011), we now suggest (Figure 8) that it is the combination of epigenetic remodeling and replication timing state that leads to the increase of chromosomal rearrangements in late-replication. We hypothesize that bias towards *cis* or *trans* chromosomal rearrangement is related to the spatial and temporal positioning of early-replicating compared to late-replicating loci (Figure 8). That is the chance of a translocation occurring is proportional to the degree of interaction between two loci and interactions occur more frequently between loci within the same chromosome territory or between adjacent territories than between distant territories (Branco and Pombo, 2006; Meaburn et al., 2007; Yaffe and Tanay, 2011). As late-replicating loci are more self-contained (Lieberman-Aiden et al., 2009), we propose that DNA breaks are more likely to involve *cis* rearrangements. In contrast as early-replicating loci are more interactive (Brown et al., 2008; Lieberman-Aiden et al., 2009), and therefore DNA breaks occurring in regions replicating in early S-phase are more likely to result in *trans* translocations.

**Figure 8:**
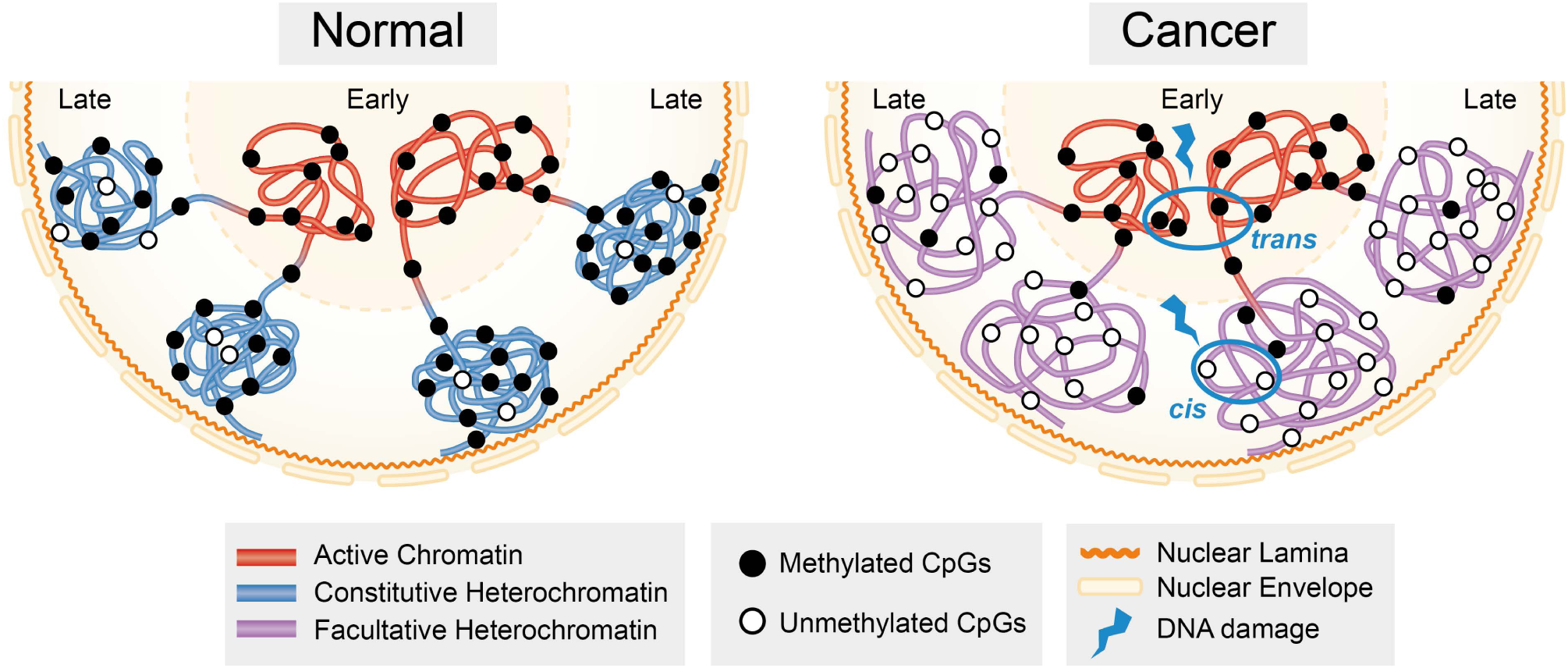
Late replication timing is more sensitive to genetic and epigenetic damage. Replication timing is spatially organised within the nucleus. Transcriptionally active early-replicating loci are often close in proximity with each other in transcriptional hubs towards the nuclear centre. Transcriptionally inactive late-replicating loci are typically heterochromatin, condensed and localised to the nuclear periphery and nuclear lamina. In normal cells, early-replicating loci are generally marked by active chromatin marks, whereas late-replicating loci are condensed and generally marked by constitutive heterochromatin (H3K9me3, 5mC). In cancer cells, early-replicating loci remain transcriptionally active, however, late-replicating loci lose constitutive heterochromatin and gain facultative heterochromatin (H3K27me3). This leads to cancer late-replicating loci becoming more open, and thus more susceptible to DNA damage that leads to chromosomal rearrangements. In relation to the *cis/trans* pattern of breakpoints, we hypothesise that if a DNA break occurs in late replication due to increased openness of chromatin; it is more likely to be repaired in *cis* as that is what is closer in nuclear space. In contrast, if a break occurred in early replication, there are more opportunities for that break to be repaired in *trans* due to the increased potential for interchromosomal interactions within structures like transcriptional hubs.

Our model is supported by studies that show copy number aberration breakpoints generally have the same replication timing and interact long-range (De and Michor, 2011), and that translocation partners are required to be within the same spatial area, transcription (Ugarte et al., 2015) or replication factory (Coll-Bastus et al., 2015), before translocation can occur. A pertinent example for prostate cancer is the TMPRSS2-ERG gene fusion that occurs in ~50-80% of all prostate cancer (Tomlins et al., 2005). Studies show that nuclear spatial proximity between TMPRSS2 and ERG is a determining factor of fusion frequency (Mani et al., 2009). Furthermore, spatial proximity can be induced through activation of the genes by the androgen receptor (AR) under testosterone (DHT) treatment, which works to decrease spatial proximity by targeting TMPRSS2 and ERG to the same replication factory (Bastus et al., 2010; Coll-Bastus et al., 2015; Mani et al., 2009). In conclusion, our combinatorial data analysis reveals a new paradigm, namely that differences in epigenetic deregulation between early and late replication shape the cancer mutational landscape and underpin long-range epigenetic deregulation of the cancer genome.

## Author contributions

S.J.C. conceived and coordinated the overall study. S.A.B., Q.D., S.S.N., A.K. W.Q and J.Z.S performed the experiments. S.A.B., Q.D. and S.J.C. interpreted data and wrote the manuscript. N.J.A. processed the Repli-Seq data, contributed to data interpretation and writing of the manuscript. C.E.C. helped with designing and performing Repli-Seq experiments. C.M.G. processed the ChIP-seq and RNA-seq data. P.L.L., E.Z. and C.S. contributed to interpretation of data. All authors read and approved the final manuscript.

## Methods

### Cell culture

LNCaP prostate cancer cells and PrEC normal prostate epithelial cells were cultured as described previously (Coolen et al., 2010).

### Repli-Seq

#### Generating BrdU-labeled ssDNA

BrdU-labeled DNA was generated as previously described (Hansen et al., 2010; Ryba et al., 2011). Briefly, cells were labeled with BrdU (50uM, Sigma, #B5002) for two hours. Labeled cells were sorted into 6 fractions across the cell cycle (G1b, S1, S2, S3, S4, G2M) as per protocol on the FACSAriaII. DNA extraction and BrdU-labeled DNA immunoprecipitation were performed with anti-BrdU antibody (40uL of 25ug/mL, BD Pharmingen, #555627). Validation of BrdU immunoprecipitation was carried out using qPCR on known Early (*BMP1*) and Late (*DPPA2*) loci (primers from Ryba *et. al.* (2011), Figures S1A and S1B).

#### Generating and sequencing Repli-Seq libraries

ssDNA was reconstituted for the complementary DNA strand using Klenow extension with random hexamers (Random Primers DNA Labeling System, Invitrogen #18187-013) using 10 ng of ssDNA input and a 2hr incubation. Reconstituted dsDNA was rechecked for enrichment of known Early and Late loci using qPCR (Figure S1C). Klenow-treated products were sonicated using a Sonifer 250 probe sonicator. 15uL of Klenow-treated dsDNA was sent to University of Southern California Epigenome Centre Data Production Facility for 50 bp single-end sequencing on the Illumina HiSeq 2000. The DNA amounts and full sequencing details can be found in Table S1.

#### Data processing

Replication Timing weighted average (WA) scores were calculated according to Hansen *et. al.* (2010) with slight modifications. To avoid bias from duplications or repeats, read densities were calculated in 150 bp intervals and intervals were excluded from further analysis if they contained greater than 20 reads per 150 bp window. Read densities were calculated in 50 kb sliding windows at 1 kb intervals across the remaining genomic regions, excluding chrY and chrM. Read densities were normalised to reads/counts per million and 50kb windows with low coverage (5 reads/counts per million) were removed. To account for variation in sequencing coverage, mapability and cell-type specific copy-number variations, the remaining 50 kb window reads/counts per million values in all 6 fractions of a given sample were converted to a percentage of total signal at each 1kb locus called Percent-normalised Density Values (PNDV) (Figure S1D). The PNDV value represents the percentage of replication occurring within a particular timing fraction at a given 1 kb locus. PNDV values were then converted into a single replication timing weighted average score per 1kb loci using the following formula: Weighted Average = (0.917*G1)+(0.750*S1)+(0.583*S2)+(0.417*S3)+(0.250*S4)+(0*G2). The formula for this transformation was obtained from the ENCODE method for “Replication Timing by Repli-Seq”. WA values represent the time of replication, where a higher WA was indicative of an earlier time of replication.

#### Quality control and data finalization

Repli-Seq was performed in duplicate for each cell line. Each sample was processed independently up to and including calculation of the WA value. WA values for replicates of LNCaP and PrEC are highly correlated (r^2^ values >0.99) (Figure S1E). Duplicate WA values per cell line were then averaged and used for downstream analysis. The distributions of WA scores were comparable to the WA distributions in other normal and cancer cell Repli-Seq datasets (Figure S1F).

#### Replication timing thresholds

Early- and late-replicating regions were defined as those regions in the top and bottom 10% of WA scores in both cell lines. This definition gives upper and lower limits of 75 and 20 respectively for PrEC and LNCaP (i.e. early regions have WA > 75 and late regions WA < 20). WA thresholds for a change in timing were defined by the following process: differences in WA between replicates of the same cell line are representative of random noise and therefore can be used as an empirical null distribution for the hypothesis test that the WA difference between LNCaP and PrEC is equal to zero. The maximum observed difference, in our data, between replicates is |∆WA| = 23 (Figure S1G). To be conservative, we chose a cutoff of |∆WA| > 25 as a value that occurs infrequently (or never) by chance under the null. Differences in WA that are larger than +/− 25 ∆WA are therefore considered to show a robust change in replication timing. To identify blocks of loci with changed replication timing, we merged all loci within 50 kb that had |∆WA| > 25. For MCF7, the top 10% of early- and late-replicating loci gives upper and lower WA limits of 73 and 17.

### Whole genome bisulphite sequencing and DMR calling

PrEC and LNCaP WGBS libraries were performed and processed as previously described (Pidsley et al., 2016). Differentially methylated regions (DMRs) were called from PrEC and LNCaP WGBS data using the package MethPipe (Song et al., 2013). The MCF7 WGBS library was performed using the CEGX TrueMethyl Whole-Genome kit (v2.1).

### RNA-seq

RNA-seq experiments were performed as previously described (Taberlay et al., 2016).

### ChIP-seq assay and processing

ChIP assays were performed as previously described (Bert et al., 2013; Coolen et al., 2010; Taberlay et al., 2014; Valdes-Mora et al., 2012) for the following histone marks, H3K4me3 (Abcam, #ab8580), H3K4me1 (Active Motif, #39297), H3K36me3 (Abcam, #ab9050), H3K27ac (Active Motif, #39133), H2AZac (Abcam, #ab18262), H3K9ac (Millipore, #06-599) and H3K27me3 (Millipore, #07-449). ChIP of H3K9me3 (Diagenode, #C15500003) was performed as previously described (Hattori et al., 2013). We performed lamin ChIP assays in PrEC and LNCaP as previously described (Lund et al., 2015) for both Lamin B1 (Abcam, #ab16048) and Lamin A/C (Santa Cruz, #sc7292). Libraries were prepared with the Illumina TruSeq Chip Library Prep Kit and sequenced on an Illumina HiSeq2500. Sequencing data was processed as previously described (Bert et al., 2013; Taberlay et al., 2014). All histone ChIP-seq peaks were called using peakranger (Feng et al., 2011). Broad domains of lamins (lamina-associated domains LADs), H3K9me3 and H3K27me3 were called using the Enriched Domain Detector (EDD) for identification of wide genomic enrichment domains (Lund et al., 2014).

### DNase1 hypersensitivity assay

Cells (7x10^6^ per sample) were scraped, centrifuged and washed with PBS. Cell pellets were resuspended in nuclear extraction buffer (10mM Tris-HCl pH7.4, 12.5mM NaCl, 3mM MgCl, 0.1mM EDTA, 0.5% IGEPAL) and dounced until nuclei were visible under light microscope with 0.4% Trypan Blue staining. DNase1 (Roche, #04716728001) was added to nuclei pellets of LNCaP (24U) and PrEC (12U) and incubated at 37°C for 15 minutes. DNaseI reactions were terminated by the addition of 36mM EDTA and Proteinase K was added before incubating at 55°C overnight. DNA was purified by phenol-chloroform extraction and ethanol precipitation. Samples were separated using electrophoresis on a 1% agarose gel. 100-300bp sections were excised and purified using the QIAquick Gel Extraction kit. Libraries were prepared with the Illumina TruSeq Chip Library Prep Kit and sequenced on an Illumina HiSeq2000.

### Quantification of epigenetic marks over replication timing loci

We defined chromatin mark occupancy and methylation averages for the 1kb wide blocks produced in the Repli-Seq data processing. Chromatin-mark-occupied 1kb blocks were defined as any 1kb block that overlapped a called ChIP-seq peak. Methylation values were averaged over the same 1kb blocks using the overlapMeans function within R package aaRon (<u>https://github.com/astatham/aaRon.git</u>). Hypomethylation was defined as a loss of > 0.2 between PrEC and LNCaP, and hypermethylation was defined as a gain of > 0.2 between PrEC and LNCaP.

### Quantification of epigenetic marks over promoters

Promoters were defined as +/− 1000bp around the start of genes from the GENCODE 19 reference transcriptome. CpG-island promoters were defined as any promoter (2 kb) that overlapped with a CpG-island. Early- and late-replicating genes were defined by calculating the average WA score over +/− 1000bp around the TSS. The genes with average WA scores > 75 or < 20 were labelled as early and late-replicating, respectively. Promoter-centric ChIP-seq enrichment for H3K4me3 and H3K27me3 was calculated by counting the number of reads within the +/-1000bp promoter region using the Repitools R package (Statham et al., 2010). Fold changes (logFC) were computed as the log2 ratio of normalised counts per promoter using the edgeR R package (Robinson et al., 2010). Methylation values were averaged over promoter regions using the overlapMeans function within R package aaRon (<u>https://github.com/astatham/aaRon.git</u>).

### CpG-island annotation

The CpG-islands are from Gardiner-Garden and Frommer (1987), downloaded from UCSC. We annotated the CpG-islands to promoters based on overlaps to promoter regions (+-1000bp) of TSS from GENCODE 19 genes. Intragenic CpG-islands were ones that overlapped the coordinates of the whole gene but did not overlap the promoter region. The rest of the CpG-islands were considered intergenic. For Figure 3E, ‘percentage of potential CpGi’ refers to the percentage of CpGi-islands with an appropriate methylation level in PrEC to show hyper- or hypomethylation. For hypomethylation, we only considered CpG-islands with methylation values of at least 0.2 in PrEC, and for hypermethylation, we only considered CpG-islands with methylation values below 0.8 in PrEC.

### Statistical tests

For genomic interval overlaps and genomic rearrangement overlaps, we modified the LOLA (Sheffield and Bock, 2016) package to perform a two-sided log odds ratio test and reports significance using “BH” FDR value. Differences between percentages of epigenetic elements in Early or Late timing were assessed using the two-sample test of equal or given proportions (prop.test in R). The Student’s T-test was used to test for significant difference between two groups of logFC values as produced from edgeR processing.

### Creating a randomized set of LRES and LREA domains for testing statistical association of domains to early or late replication

Random genomic regions were generated in a three stage process: first, a chromosome was selected at random, second, the start point of the region was randomly generated from a uniform distribution between 1 and the length of the chromosome, and last, the length of the region to be generated by sampling at random from the known lengths of LREA or LRES regions. If the random region generated did not fit on the chromosome, it was discarded and the process repeated. In this way, we generated 1000 regions across the genome that were distributed in length similarly to the LREA and LRES regions. We computed the WA for each random region and compared this empirical null distribution to the distribution of WA values for LREA and LRES using the Mann-Whitney-Wilcoxon test.

### Profile plots

We used genomation (Akalin et al., 2015) to calculate average WA scores over regions of interest, which were divided into 50 bins per region. We then used ggplot (Wickham, 2016) to plot the average WA scores across all regions for each bin with standard error and confidence intervals.

### Lamina boundary heatmaps

We performed the below separately for LMNA and LMNB1 LADs. Only LADs that overlapped between PrEC and LNCaP were used. We calculated the bp distance from the PrEC LAD 5’ or 3’ boundary to the nearest LNCaP LAD 5’ or 3’ boundary, respectively. Negative and positive distances denote that the LNCaP boundary is respectively upstream or downstream of the PrEC boundary. Heatmaps of WA values are centred on PrEC LAD boundaries, and are ordered by decreasing distance to the nearest LNCaP boundary.

### Conserved replication timing alterations in cancer

WA values from all publically available Repli-Seq datasets and our datasets were scaled prior to performing PCA and hierarchical clustering. PCA was performed on 1kb loci that were present in all datasets. Hierarchical clustering was performed using the hclust function in R with the ward.D2 method. To find conserved regions of changed timing in cancer (LNCaP, MCF7, SK-N-SH, HepG2, K562, HelaS3) compared to all other Repli-Seq datasets (See Figure 6A), WA values were quantile-normalized and scaled before using limma (Ritchie et al., 2015) to find regions of difference. We filtered for regions that were significant (fdr-corrected p-value < 0.05) with a logFC >= 1. These regions were merged if they were within 50kb of each other to give the final set of Earlier Cancer Domains (ECDs) and Late Cancer Domains (LCDs). We further filtered ECDs and LCDs through overlap with high and low replication timing variation regions. These high/low variation regions were defined as the top and bottom 20% of 1kb loci based on scores outputted by the ‘var’ function in R. Repli-Seq scores per loci from all was used as input for ‘var’ to calculate a score per loci. We further separated ECDs and LCDs into ones that associate with replication timing of ESC. We classified a region as associated with ESC if the difference between the averaged replication timing score of cancer to ESC was less than 0.5 for ECDs or more than −0.5 for LCDs (quantile-normalised and scaled WA scores).

### Gene set enrichment analysis for genes in ECDs and LCDs

We were unable to call significant terms from GSEA with the stringent cutoffs initially used to define ECD and LCDs (logFC >= 1, See above) due to the restricted number of genes obtained. We relaxed our domain calling cutoff to logFC > 0 to obtain a less stringently defined but larger list of genes for GSEA analysis. We used a hyper-geometric test to scan the MolSigDB v6.0 (Liberzon et al., 2015; Subramanian et al., 2005) for gene sets with statistically significant overlap with genes found within ECDs and LCDs. More specifically, we computed the overlap between the MolSigDB gene sets and our set of genes and compared what would be expected by chance if equivalent number of genes were drawn uniformly at random from the background set of genes. We report statistically significant enrichments with FDR < 0.05 in Figure S7.

### Public datasets

For prostate cancer breakpoints, we used the Baca *et al.* (2013) (Baca et al., 2013), Berger *et al.* (2011) (Berger et al., 2011), and Robinson *et al*. (2015) (Robinson et al., 2015) datasets. Publicly available Repli-Seq datasets used in this study were downloaded from ENCODE data portal https://www.encodeproject.org/matrix/?type=Experiment&assay_title=Repli-seq. These datasets were created by the University of Washington ENCODE group (Hiratani et al., 2010; Thurman et al., 2007). Raw sequence data was downloaded from UCSC, mapped to hg19 and processed in the same manner as our own Repli-Seq data as described above. HMEC and MCF7 ChIP-seq data was downloaded from ENCODE. HMEC ChIP-seq is from https://www.encodeproject.org/reference-epigenomes/ENCSR044VTN/ and MCF7 ChIP-seq is from https://www.encodeproject.org/reference-epigenomes/ENCSR460EGF/. HMEC WGBS was downloaded from GSE29127 and processed in-house.

### Data and GEOs

Raw and processed Repli-Seq and ChIP-seq data from this study have been submitted to the NCBI Gene Expression Omnibus (GEO; http://www.ncbi.nlm.nih.gov/geo/) under accession number GSE98732. RNA-seq data from Taberlay *et al.* (Taberlay et al., 2016) are available under GEO accession number GSE73784. ChIP-seq data from Valdes-Mora *et al*. (Valdes-Mora et al., 2012), Bert *et al.* (Bert et al., 2013), Taberlay *et al*. (Taberlay et al., 2014) and Taberlay *et al.* (Taberlay et al., 2016) are available under GEO accession numbers GSE25914, GSE38685, GSE57498, GSE73785, respectively. WGBS data from Pidsley *et al.* (Pidsley et al., 2016) are available under GEO accession number GSE86833. A summary of the new and existing data used in this manuscript for PrEC and LNCaP can be found in Table S4.

## Supplemental Figure Legends

**Supplementary Figure 1: Validation of Repli-Seq datasets and determining replication timing difference thresholds.**

(**A**) Validation of correct S-phase sorting. qPCR of known early (BMP1) and known late (DPPA2) loci was carried out on BrdU-labelled DNA extracted from sorted PrEC and LNCaP samples. (**B**) Relative levels of BMP1 to DPPA2 loci were used as a measure of early to late replication. The y-axis uses a log_10_ scale. (**C**) qPCR was repeated on the same samples following Klenow dsDNA synthesis. (**D**) Raw read densities across all S-phase fractions are normalised to give a Percent Normalised Density Values (PNDV) per 1kb locus for each fraction. Red arrows indicate regions of replication initiation and the blue arrow indicates a region of replication termination. PNDV values of all 6 fractions are used to calculate a single Weighted Average (WA) score per locus. Higher WA values indicate early replication timing and lower WA values indicate late replication timing. (**E**) Replicates of each cell line show high correlation and r^2^ scores. (**F**) The distribution of PrEC (green) and LNCaP (red) WA values are comparable to the WA distributions of twelve ENCODE Repli-Seq datasets (pink). (**G**) The distribution of WA differences between replicates (left) is compared to the distribution of WA differences between PrEC and LNCaP (right). Dotted lines indicate a |∆WA| > 25. Values that fall outside this range (red) signify loci that have changed replication timing.

**Supplementary Figure 2: Replication timing in the normal and cancer prostate genome.**

Repli-Seq derived PNDV values of all cell fractions are plotted for each chromosome (hg18). The name and size of each chromosome is shown above the respective chromosomal ideogram. Green values represent PrEC and red values represent LNCaP. Domains that replicate *earlier* in LNCaP are highlighted in blue; domains that replicate *later* in LNCaP are highlighted in magenta.

**Supplementary Figure 3: Relationship between chromatin marks and replication timing.**

Percentage occupancy of chromatin marks for 1kb loci across replication timing for active (**A**) and repressive (**B**) marks in PrEC and LNCaP. Each mark is compared against the percentage of genome occupancy for that mark. The same was done for HMEC and MCF7 using MCF7 replication timing (**C**). Log odds ratio tests for overlap between loci that change in replication timing and loci that change in active (**D**) or repressive (**E**) chromatin mark occupancy between PrEC and LNCaP. Positive values (RHS) indicate association between the difference in chromatin mark occupancy and the change in timing. Negative values (LHS) indicated disassociation between the difference in chromatin mark occupancy and the change in timing. Asterisks indicate significant associations (FDR < 0.05).

**Supplementary Figure 4: Relationship between DNA methylation and promoter-CpG-island genes with replication timing.**

(**A**) DNA methylation (WGBS) density distributions for early (blue) and late (red) loci for HMEC and MCF7. Adjacent are scatterplots of DNA methylation in relation to replication timing (WA) for all measured 1kb loci in HMEC and MCF7. Blue dashed line indicates early (WA > 73) and red dashed line indicates late (WA < 17). (**B**) DNA methylation of LNCaP across LNCaP replication timing percentiles, separated by genomic elements. (**C**) Representative examples of late-replicating regions in both PrEC and LNCaP (shaded) that become hypomethylated in LNCaP. (**D**) Expression (mean.TPM) of early and late genes in PrEC and LNCaP, separated by CpG-island status. (**E,F**) Scatterplots showing the relationship between change in replication timing and change in H3K27me3 and H3K4me3 read density between PrEC and LNCaP at gene promoters. Dashed lines indicate a |∆WA| > 25, outside of which indicates a change in replication timing (red points). logFC boxplots for *later* or *earlier* genes are on the left and the right of the scatterplot, respectively. *Later* or *earlier* genes are significantly changed in expression compared to genes between |∆WA| > 25 (asterisk, Student’s T-test, p < 2.2e-16).

**Supplementary Figure 5: Long-range epigenetically regulated domains in cancer change in replication timing.**

(**A**) Average plots of PrEC and LNCaP WA values over regions containing both lamin A/C and B1 LADs, that are conserved between PrEC and LNCaP, lost in LNCaP or gained in LNCaP. (**B**) Heatmap of PrEC and LNCaP WA values over the boundary of PrEC lamin A/C LADs, ordered by degree of lamin A/C extension (upstream of 5’ and downstream of 3’) and loss (between 5’ and 3’) in LNCaP. Black lines down the centre of heatmaps represent PrEC LAD boundaries. White lines indicate the lamin A/C boundary in LNCaP. Scale for WA is from late (red) to early (blue). (**C**) The top ten most common combinations of changes in DNA methylation, lamina association, and heterochromatin within LNCaP late-replicating loci. The table below illustrates which combination of changes has occurred. Comparing LNCaP to PrEC, “-” indicates loss of that mark, “+” indicates gain of that mark, “=” indicates that the mark is present and not changed, and “0” indicates that the mark is not present in either cell line. (**D**) A representative example of a late-replicating region in LNCaP showing maintained LADs with coordinate DNA hypomethylation and H3K27me3 gain, without presence of H3K9me3. (**E**) Examples of domains of replication timing change overlapping LRES and LREA regions. PrEC and LNCaP data are respectively represented by the colors green and red. Blue shading indicates an *earlier* domain. Pink shading indicates a *later* domain.

**Supplementary Figure 6: Replication timing of cancer compared to other cell types in Earlier Cancer Domains and Later Cancer Domains.**

Averaged replication timing scores for cancer (HEPG2, K562, MCF7, SKNSH, LNCaP, HELAS3), embryonic stem cell (BG02-ESC), lymphoblastoids (GM06990, GM12801, GM12812, GM12813, GM12878) and normal (BJ_Rep1, BJ_Rep2, IMR90, HUVEC, NHEK, PrEC) across ECDs and LCDs are represented by the dot along each vertical parallel coordinate for that cell group. Each line represents a single domain and connects the averaged scores (dots) of the 4 cell groups for that domain. ‘Variable’ indicates domains that overlap with regions of high variation between cell types (See Methods). ‘ESC’ indicates domains where cancer replication timing corresponds to ESC replication timing (See Methods). ECDs show consistently higher replication timing scores in cancer compared to lymphoblastoid and normal, and higher than ESC in domains that do not associate with ESC timing (‘Variable, not ESC’). LCDs show consistently lower replication timing scores in cancer compared to lymphoblastoid and normal, and lower than ESCs in domains that do not associate with ESC timing (‘Variable, not ESC’). The timing scores shown here are after quantile normalisation and scaling (See Methods).

**Supplementary Figure 7: Significant GSEA terms for genes found within ECDs or LCDs.**

ECD and LCD genes from domains called with a logFC > 0 cutoff were tested against GSEA MolSigDB v6.0 gene sets. Only gene sets with significant terms are shown here. We used a hyper-geometric test to identify statistically significant enrichment (p-value < 0.05, bonferroni corrected) of gene sets with ECD and LCD genes. We show the top 10 terms ranked by p-value. We also show fold enrichment of ECD/LCD genes for each significant gene set term. Dotted line on the −log10p-value axis is the significance cutoff. Dotted line on the Fold Enrichment axis is where fold enrichment = 1.

**Supplementary Figure 8: Examples of replication timing changes relating to long-range domains and gene fusions in prostate cancer**

(**A**) Replication timing (PrEC WA) distributions for chromosomal rearrangements divided into tandem duplications, inversions, deletions and long-range cis rearrangements. Solid lines are distributions for Baca *et. al.* (2013) and dashed lines are distributions for Berger *et. al.* (2011). (**B**) Replication timing differences within PrEC and LNCaP (top and middle panel) between the 5’ and the 3’ gene fusion breakpoints described from clinical prostate cancer samples (Robinson et al., 2015), and replication timing differences between PrEC and LNCaP of the same breakpoint (bottom panel). Distributions of breakpoint timing differences are compared to genome wide timing differences between PrEC and LNCaP (dotted line). (**C**) Replication timing change across the ERG locus.

**Supplementary Table 1: Sequencing metrics and QC for RepliSeq.**

PrEC and LNCaP were sorted into 6 fractions in duplicate. DNA (ng) refers to the amount of DNA sent for sequencing at the USC Epigenome Centre. ‘Total reads’ refers to the number of raw reads obtained for each sample; ‘unique reads’ refers to the final number that were suitable for further analyses.

Supplementary Table 2: List of genes found in ECDs and LCDs.

Supplementary Table 3: Prostate cancer gene fusions and replication timing. Related to Figure S8.

Supplementary Table 4: A summary of datasets generated and used in this study.

## Acknowledgements

We thank members of the Clark Laboratory for helpful discussions and careful reading of the manuscript. In particular, we thank Madhavi Maddugoda for help with figures. S.J.C. is a National Health and Medical Research Council (NHMRC) Senior Principal Research Fellow #1063559. This work was supported by NHMRC Project Grants #1011447 and #1051757 and APCRC-NSW. The authors declare no conflicts of interest.

